# A Bioluminescent Reporter for Antibacterial Defence Induction in *Coprinopsis cinerea*

**DOI:** 10.64898/2026.08.11.743940

**Authors:** Alessandri Emma, Welman Judyta, Lohmann Luca, Künzler Markus

**Affiliations:** ETH Zürich, Department of Biology, Institute of Microbiology, Switzerland; Universität Zürich, Department of Biochemistry, Switzerland

## Abstract

The coprophilous agaricomycete *Coprinopsis cinerea* is a model organism for antagonistic fungal-bacterial interactions. Previous studies showed that *C. cinerea* responds to antagonistic bacteria with strong induction of a set of genes encoding secreted antibacterial molecules. However, little is known about the elicitors of this response. Key open questions in this respect include whether individual antibacterial defence genes are induced by different bacteria and/or by specific bacterial soluble molecules.

Here, we present a new *C. cinerea* reporter system to monitor antibacterial defence induction and address related outstanding issues with minimal hands-on time. In this system, the promoter of the endogenous bacterial-induced gene *cclys1* drives the expression of *cnluc*, which encodes a secreted variant of the deep-sea shrimp luciferase Nluc. We show that cNluc allows to detect and quantify *cclys1* induction by measuring luminescence directly in the culture medium of reporter strain colonies. Building on these features, we successfully leveraged the inducible cNluc reporter strain for the development of a novel 96-well plate assay that allows the high-throughput screening of antibacterial defence elicitors. As cNluc can be subject to degradation by secreted proteases of fungal or bacterial origin in the culture medium, we coupled this assay to confirmatory qRT-PCR. Testing this set-up by confronting the reporter strain with several different bacteria revealed that *cclys1* induction occurs independently of the bacterial ecological niche. Based on these results, we also recommend qRT-PCR exclusively for validation of negative results. We conclude that cNluc offers significant advantages over cytoplasmic reporter proteins, especially for preliminary rapid screening of multiple conditions.

## Introduction

Fungi share their ecological niches with bacteria, which leads to cross-kingdom competition over limited resources such as space and nutrients. In order to strengthen their competitiveness, fungi have evolved different strategies, which involve the production of antibacterial molecules (i.e. chemical defence). The production of these molecules can be either constitutive or induced in response to antagonistic bacteria [1, 2].

The coprophilous agaricomycete *Coprinopsis cinerea* deploys both constitutive and inducible chemical defence layers against bacteria and is thus a model organism for the study of antagonistic fungal-bacterial interactions. Constitutively produced antibacterial molecules include two intracellular lactonase enzymes, which hydrolyze quorum sensing signalling molecules produced by Gram-negative bacteria [3]. Another notable example is copsin, an antimicrobial peptide (AMP) of the cysteine-stabilized α/β-defensin class, which is particularly active against Gram-positive bacteria [4, 5].

The inducible layer of antibacterial defence in *C. cinerea* has been explored in more recent studies. Kombrink et al. [6] found that co-cultivation with either *Bacillus subtilis* or *Escherichia coli* results in the significant overexpression of an overlapping set of genes. These genes mostly encode secreted antibacterial products, including a paralog of copsin (*cpp2*) and bacterial cell-wall targeting lysozymes (*lys1* and *lys2*). Lysozyme-encoding genes are among the most highly induced upon either treatment, together with the biosynthetic cluster of lagopodin B, an antimicrobial secondary metabolite [7]. Interestingly, the cell-free supernatant (CFS) of *B. subtilis* is sufficient to induce these genes [7]. This observation suggests that *C. cinerea* may be able to also sense and respond to bacterial-derived soluble molecules. To date, the nature and specificity of these elicitors, particularly with regard to the induction of individual antibacterial defence genes, remain elusive. It is also unclear if these defence genes are induced by bacteria other than the model organisms *B. subtilis* and *E. coli*, whose ecological niches overlap with *C. cinerea*.

To answer the above questions, we originally developed a *C. cinerea* reporter strain by fusing the promoter of *lys1* (*cclys1*p) to the coding region of the cytoplasmic red-fluorescent protein dTomato. Production of dTomato faithfully reports *lys1* expression, but monitoring intracellular fluorescence requires cultivation of the reporter strain in custom-made polydimethylsiloxane (PDMS) microchannel devices [7]. PDMS devices are expensive and limited in supply. For this reason, we chose to still rely on our traditional challenge assays, which are instead carried out in Petri dishes by growing *C. cinerea* on liquid minimal medium, supported by a bed of glass beads (i.e. semi-liquid conditions) [5, 6]. In this set-up, dTomato production is assessed by immunoblotting the whole mycelium protein extract (WMPE) of induced fungal colonies with anti-dTomato antibodies [7]. These assays are laborious, destructive (i.e., they require protein extraction), and semi-quantitative at best.

Here, we addressed the above limitations with a new *C. cinerea* reporter strain, in which *cclys1*p drives the expression of a gene encoding a secreted variant of the luciferase Nanoluc (Nluc), hereafter referred to as ‘cNluc’. To control for cNluc activity and stability under our experimental conditions, we also generated another reporter strain, which secretes cNluc constitutively. This was achieved by fusing *cnluc* to the strong constitutive promoter of the glyceraldehyde-3-phosphate dehydrogenase encoding-gene (*gpdII*) from *Agaricus bisporus* (*abgpdII*p) [8].

We validated the inducible cNluc reporter strain in co-cultivation with *B. subtilis* and *E. coli* using the semi-liquid set-up. Our results show that the cNluc system 1) does not require protein extraction, which minimizes hands-on time, 2) allows quantification of *lys1* induction levels, and 3) is highly sensitive. These three features made it possible to adapt the challenge assays to 96-well plates, which in turn allows the high-throughput screening of candidate antibacterial defence elicitors. As a first step in this direction, we tested the new set-up by confronting the inducible cNluc reporter strain with bacteria from various ecological niches. As a result, we were able to clarify that *lys1* is also induced by bacteria that do not share the same ecological niche as *C. cinerea*. While cNuc enzymatic activity may be reduced or lost under specific conditions, we demonstrate that reporting gene expression with cNluc still offers considerable advantages over other approaches, including its suitability for high-throughput screenings.

## Materials and Methods

### Strains and cultivation conditions

Fungal and bacterial strains used in this study are summarized in Table S1.

*C. cinerea* was cultivated on solid yeast-extract malt-extract glucose (YMG) medium at 37°C in aerated, dark and humid boxes, unless stated otherwise. *Saccharomyces cerevisiae* W303 *MAT*a was cultivated at 28 °C on yeast-extract peptone dextrose (YPD) medium and selected on synthetic complete dextrose without uracil (SD Ura-) medium when used for homologous recombination. *E. coli* DH5α was used for cloning and maintenance of plasmids. Preparation of competent cells and transformations were carried out as previously described [9]. Selection of positive colonies was carried out at 37°C on Luria Bertani (LB) medium supplemented with 100 μg ml^−1^ ampicillin. All bacterial strains used in co-inoculation experiments were pre-grown at 28 °C on LB agar plates.

### Construction of cNluc-secreting *C. cinerea* reporter strains

The coding sequence of the luciferase Nanoluc (Nluc) was retrieved from plasmid pNL1.1 (Promega, Madison, WI) and codon-optimized for expression in *C. cinerea* with ‘Dicodon optimizer’ [10]. The coding region was also modified to C-terminally tag the protein with the 1D4 epitope of bovine rhodopsin (Rho1D4). To direct the protein to the culture medium, a sequence coding for the secretion signal of *C. cinerea lys1* (*cclys1*, CC1G_03076T0), as defined by SignalP 6.0 [11], was added to the 5’end of the Nluc-Rho1D4 coding sequence. The *Nluc* coding sequence with the above modifications (‘*cnluc*’, Fig. S1) was then chemically synthesized and inserted into a pTwist Amp High Copy vector (Twist Biosciences, San Francisco, CA).

For expression in *C. cinerea*, the *cnluc* coding region was fused to either a constitutive or bacteria-inducible promoter by homologous recombination into existing expression vectors. For constitutive expression, *cnluc* was recombined into plasmid pMA412 under control of the *gpdII* promoter from *Agaricus bisporus* (*abgpdII*) (Fig. S2A) [8]. For expression upon bacterial induction, *cnluc* was introduced into plasmid pMA1070 under control of the *cclys1* promoter from *C. cinerea* (*cclys1*p) (Fig. S2B) [7]. In both constructs, *cnluc* is under the control of the 5’ UTR and terminator regions of *mnp* from *Phanerochaete chrysosporium* (*pcmnp*t). Furthermore, an intron from the *P. chrysosporium mnp* gene was introduced between the start codon and the first amino acid coding triplet of the *cnluc* coding sequence, to increase expression efficiency as previously reported [12]. The *cnluc* coding sequence was amplified from the pTwist plasmid using primers containing homology regions to either pMA412 or pMA1070. Both destination plasmids were linearized using restriction enzymes *FspAI* and *BsrGI*. Homologous recombination was carried out in *S. cerevisiae* W303 *MAT*a, as described previously [13]. Resulting plasmids were verified by DNA sequencing (Microsynth, Balgach, CH) and transformed into *C. cinerea AmutBmut* via ectopic integration, as described previously [14].

### Semi-liquid set-up for the co-cultivation of fungi and bacteria

Strains were assessed in a semi-liquid set-up as previously described [5]. Briefly, one agar plug from the edge of a *C. cinerea* colony grown at 28 °C (3 days) was inoculated in a Petri dish (55 mm diameter) onto a layer of borosilicate glass beads (5 mm diameter, Sigma-Aldrich) submerged in 5 mL liquid *C. cinerea* minimal medium (CCMM) pH 6.4. Plates were incubated at 28 °C for 2.5 days in a dark humid box prior to inoculation with bacteria. Bacteria were grown in CCMM at 37 °C to an optical density at 600 nm (OD_600_) of 0.3, pelleted, and resuspended in fresh CCMM to OD_600_= 0.6. Each fungal plate was then inoculated with 500 µL of bacterial resuspension and incubated overnight at 28 °C in the dark, humid box. Bacterial cell-free supernatant (CFS) was lyophilized, extracted with Methanol, and concentrated as previously described previously [7]. When relevant, 500 µL of concentrated CFS was inoculated into each fungal plate, followed by overnight incubation as above.

### Extraction and immunoblotting of whole mycelium protein

Fungal strains grown in semi-liquid conditions were harvested following incubation (with or without bacteria). 3 biological replicates were combined for extraction of total mycelial protein, which was performed as previously described [3]. Total protein concentrations were determined using the BCA assay. Total protein samples were boiled in Lämmli buffer and 45 µg was loaded and run out on a 12% polyacrylamide gel. The gel was transferred to a nitrocellulose membrane, which was then probed with a 1:500 dilution of the primary Rho 1D4 antibody (mouse anti-Rho recombinant antibody, Chemicon) and a 1:1000 dilution of the secondary antibody (horseradish peroxidase conjugated goat anti-mouse immunoglobulin G, Santa Cruz Biotechnology). Immunoblots were imaged in a Fusion FX7 chemiluminescence imager (Vilber Lourmat, Eberhardzell, Germany).

### Assessment of of cNluc activity and stability upon different treatments

To obtain cNluc-containing conditioned CCMM, 5 glass-bead petri dishes of 55 diameter, each containing 5 mL CCMM, were inoculated with the *abgpdII*p*-cnluc* reporter strain as described above and incubated at 28 °C in the dark, humid box. After 3 days, the conditioned CCMM was pulled, filter-sterilized, and used immediately for experiments. To investigate the impact of bacteria upon cNluc, *B. subtilis* and *E. coli* were grown in CCMM at 28 °C to an OD_600_ of 0.5, pelleted, and resuspended to OD_600_ = 1 in cNluc-containing CCMM. 500 µL aliquots of bacterial resuspensions were distributed into sterile 1.5 mL Eppendorf tubes and incubated for 12 hours at 28 °C and 300 rpm in a table-top heating block. Bovine serum albumin (BSA) (Sigma-Aldrich), concentrated LB, and the protease inhibitor cocktail (PIC) (Sigma-Aldrich) were added directly to 500 µL cNluc-containing CCMM to their respective final concentrations. Luminescence was recorded at the beginning and at the end of the incubation period.

### 96-well cultivation set-up for high-throughput screening of *cclys1*p induction

To produce oidia, the inducible cNluc reporter strain, grown for 5 days on YPD as described above, was transferred to a transparent humid box and incubated under constant white light (100% intensity) at 37°C for another 3 days. Oidia were harvested by pouring 10 mL of sterile double-distilled water (ddH_2_O) on top of the colony and gently scraping the surface with a plastic inoculation loop. This procedure was repeated twice. The oidia-containing suspension was then filtered through a 70 µm cell strainer (Greiner Bio-One) and centrifuged for 15 minutes at 4500 rpm. Pelleted oidia were resuspended in CCMM to an OD_600_ of 0.5 and 100 µL aliquots thereof were distributed into a 96-well plate (100 µL/well). The plate was incubated at 28 °C in the dark, humid box for 7 days. Bacteria were cultivated overnight in CCMM at 28 °C, diluted in CCMM to an OD_600_ of 0.5, and inoculated into the 96-well plate (on day 7) in 100 µL aliquots to a final volume of 200 µL/well. *cclys1* expression and luminescence in each well was measured following overnight incubation (on day 8) as described below.

### qRT-PCR validation of *cclys1* expression in the inducible cNluc reporter strain upon bacterial inoculation in 96-well plate

The content of each well was pulled by row, centrifuged to remove residual supernatant, frozen in liquid nitrogen, and lysed with a plastic pestle. Total RNA extraction was performed with Plant/Fungi Total RNA Purification kit (NORGENTM) according to manufacturer’s instructions, DNase treatment included. RNA quality was assessed with the Bioanalyzer 2100 (Agilent). All RNA samples had an integrity index ≥ 6 and could be used for subsequent cDNA synthesis with random hexamer primers using SuperScript™IV First-Strand Synthesis System (Thermo Fisher Scientific). qRT-PCR and data analysis were performed as previously described using *cclys1* cDNA-targeting primers listed in Table S2 [15].

### Detection and quantification of luminescence in conditioned liquid CCMM

To detect and quantify luminescence, 30 µL of conditioned CCMM per plate/well was mixed to a 30 µL solution containing buffer and the luciferase substrate furimazine in a 50:1 ratio, according to the manufacturer’s instructions (Promega, Madison, WI). All reactions were set-up simultaneously in a black, 96-well microplate (Huberlab, Aesch, CH) and incubated for 5 minutes at room temperature. Luminescence in each well was detected and quantified using a Victor3 Wallac 1420 microplate reader (PerkinElmer Life and Analytical Sciences). Measurements were normalized to sterile CCMM mixed with the buffer-furimazine solution. Pictures of luminescent 96-well plates were taken in a Fusion FX7 chemiluminescence imager (Vilber Lourmat, Eberhardzell, Germany).

## Results

### Production and secretion of functional cNluc in *C. cinerea*

The *cnluc* coding region was fused to either *cclys1*p or *abgpdII*p, resulting in the inducible and constitutive reporter construct, respectively (Fig. S1A). Both constructs were introduced into plasmids carrying the wildtype version of the para-aminobenzoic acid (PABA) synthase-encoding gene (*ccpab1*) and ectopically integrated into the genome of PABA-auxotrophic *C. cinerea* strain *AmutBmut* (see Materials and Methods section). Transformants were selected based on growth on minimal medium and confirmed by colony PCR (Fig. S3A). To obtain homokaryotic transformants, oidia from confirmed transformants were produced and germinated on selective medium to obtain single colonies. Single colonies were then screened for the strongest luminescent signal to identify the best cNluc reporter strains for the follow-up experiments. For this purpose, the homokaryotic transformants were cultivated in the same semi-liquid setup already used to investigate fungal-bacterial interactions [6, 7]. Both the constitutive (*abgpdII*p*-cnluc*) and the inducible (*cclys1*p*-cnluc*) reporter strains were grown axenically for 3 days. To induce *cclys1*p, the inducible reporter strains were inoculated with concentrated *E. coli* cell-free supernatant (CFS) on the second day as previously described [7]. Following overnight incubation, the luciferase-specific substrate furimazine was added to aliquots of conditioned culture medium in a 96-well plate. While luminescence was detectable in most aliquots, the intensity differed significantly between transformants of the same reporter construct (Fig. S3B-C). These differences are most likely due to variability in the integration of the reporter gene, in terms of frequency and genome localization, between the different transformants.

The brightest homokaryotic transformant per reporter construct were further validated under semi-liquid conditions in the presence of bacteria (Fig.1A). The constitutive reporter strain was grown axenically, while the inducible one was cultivated with or without *B. subtilis*. Expression of cNluc was assessed in both strains by immunoblotting of the WMPE with anti-Rho1D4 antibodies. We also measured luminescence in the culture medium, to confirm promoter-dependent secretion of functional cNluc. As expected, *abgpdII*p drove the expression of cNluc constitutively, while cc*lys1*p only upon bacterial induction (Fig. 1B-C).

**Figure 1.**
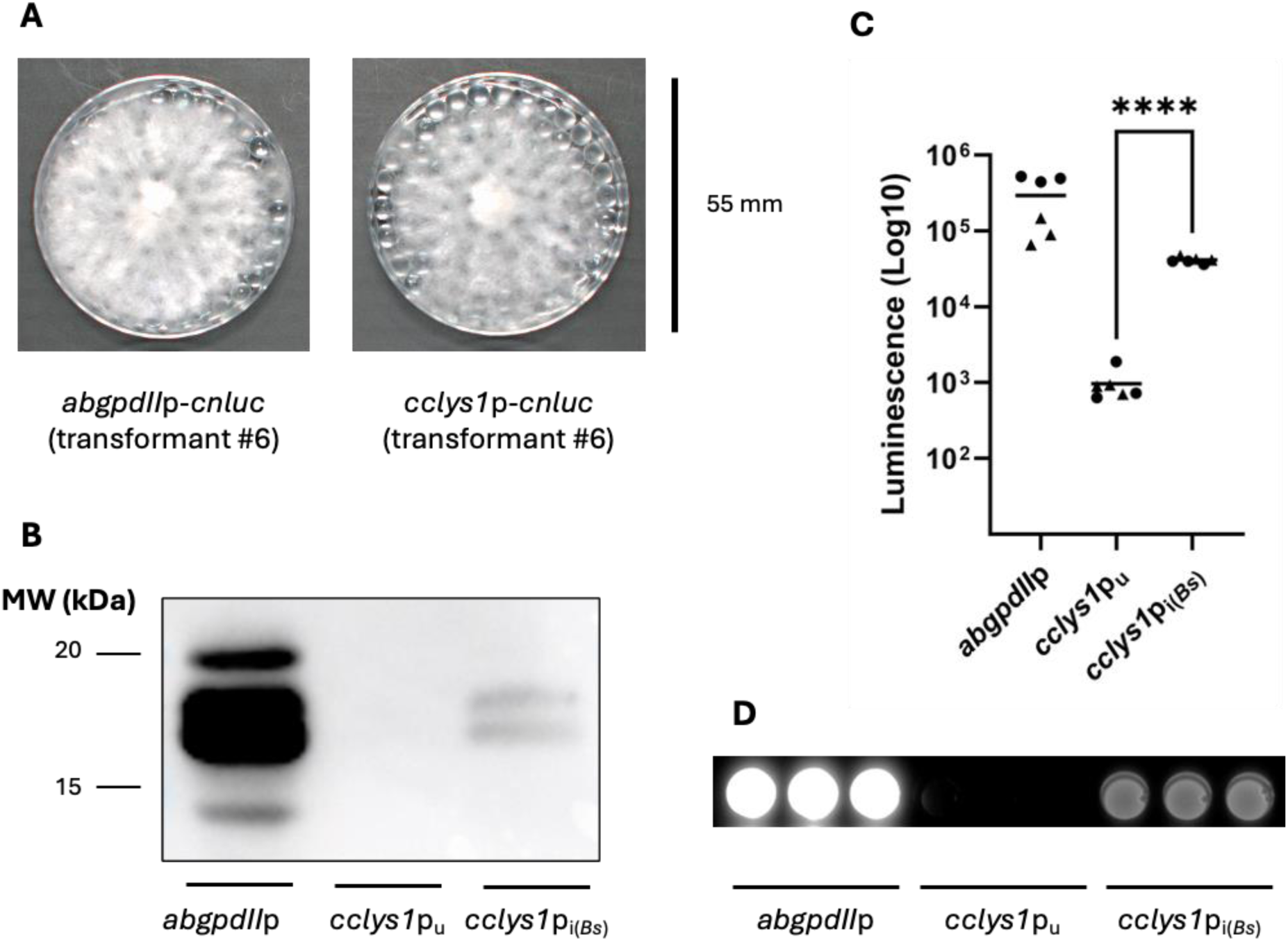
Production and secretion of active cNluc in *C. cinerea*. All data in this figure were obtained from *C. cinerea* transformant strains grown for 3 days in CCMM on borosilicate glass beads. The constitutive reporter strain (*abgpdII*p*-cnluc*) was grown axenically, the inducible reporter strain (*cclys1*p-*cnluc*) either axenically (i.e. uninduced, *cclys1*p_u_) or incubated overnight with *B. subtilis* (i.e. induced, *cclys1*p_i(*Bs*)_). **(A)** Representative images of homokaryotic *C. cinerea* transformant strains, carrying either the *abgpdII*p*-cnluc* or the *cclys1*p*-cnluc* fusion construct. These specific transformants (#6) were used in all following experiments. **(B)** Immunoblot with anti-Rho1D4 antibodies of WMPE from the constitutive or inducible reporter strain. The four bands are interpreted as follows (from top to bottom): precursor protein with signal peptide, mature protein without signal peptide, mature protein (without LYS1 signal peptide) with N-terminal deletion, mature protein (without LYS1 signal peptide) with further N-terminal deletion. **(C)** Luminescence of conditioned CCMM from cultures of constitutive or inducible reporter strain. Data from two experiments are shown. Each data point represents one biological replicate as the average of two independent measurements. Data points with the same symbol belong to the same experiment. Horizontal bars represent means. Unpaired T-test. *p < 0.05, **p < 0.01, ***p < 0.001, ****p < 0.0001. **(D)** Luminescence of 30 µL conditioned CCMM from cultures of constitutive or inducible reporter strain. Conditioned medium was distributed into wells of a black 96-well plate and imaged as described. Each well contains conditioned medium from a distinct biological replicate.

Finally, we found that both cNluc strains grow better than untransformed *AmutBmut* in liquid CCMM (Fig. S3D-E). This is due to the integration of functional PABA synthesis. While CCMM is supplemented with PABA to enable growth of untransformed *AmutBmut*, endogenous synthesis of PABA in transformed strains still appears to confer a competitive advantage.

### Validation of the inducible cNluc reporter strain for quantification of antibacterial defence induction in *C. cinerea*

The development of this new reporter system was driven by the need for a quantifiable output to detect gene induction with minimal hands-on manipulation. We show that cNluc secretion minimizes hands-on time, as its enzymatic activity can be detected in the fungal culture medium without destructing the mycelium. Moreover, the assay is quantitative as the measured luminescence is linearly proportional to the amounts of produced enzyme (Promega, Madison, WI). This allows to compare the induction strength of different treatments.

To confirm this, we compared the intensity of the luminescence produced by the inducible reporter strain upon challenge with *E. coli* or *B. subtilis* in our usual semi-liquid set-up. We had previously reported that *E. coli* induces significantly higher expression levels of *cclys1* than *B. subtilis*. This was based on comparative transcriptomic analysis [6]. In co-cultivation with the inducible reporter strain, *E. coli* reached a higher density than *B. subtilis* (Fig. S4A), but *B. subtilis* confrontation produced significantly higher levels of luminescence than *E. coli* (Fig. 2A; Fig. S4B). Since these results do not align with our previous observations, we searched for possible explanations and investigated the system more closely.

**Figure 2.**
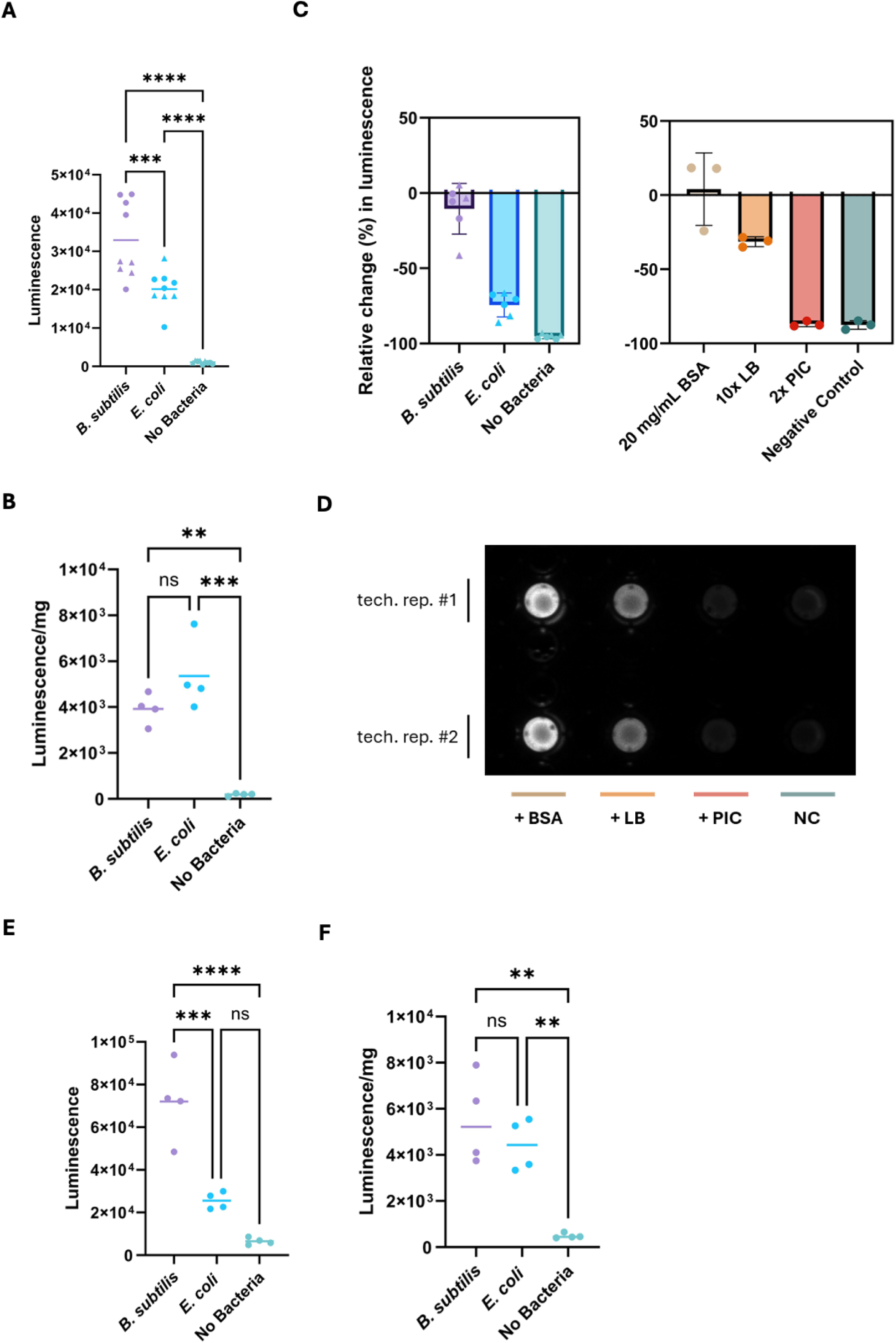
Validation of the inducible reporter strain for quantification of antibacterial defence induction. **(A)** Luminescence of conditioned CCMM from cultures of inducible reporter strain. The transformant was grown for 3 days on borosilicate glass beads and inoculated overnight with either *B. subtilis* or *E. coli*. Data from two experiments are shown. Each data point represents one biological replicate as the average of two independent measurements. Data points with the same symbol belong to the same experiment. Horizontal bars represent means. **(B)** Data from panel A (one experiment) normalized by fungal colony dry weight (mg). Horizontal bars represent means. **(C)** Relative change (%) in luminescence after 12 hrs in sterile, cNluc-containing CCMM following different treatments: addition of *B. subtilis* or *E. coli* (data from two experiments, each represented by a different symbol); bovine serum albumin (BSA), 10x-concentrated LB medium (10x LB), or 2x-concentrated protease inhibitor cocktail (2x PIC) (data from one experiment). Each treatment was performed in technical triplicates and each data point corresponds to the luminescence measured in one replicate. The bar charts also show means and standard deviations. **(D)** Representative picture of residual luminescence of the same cNluc-containing CCMM 12 hours after addition of either 20 mg/mL bovine serum albumin (BSA), 10x-concentrated LB medium, or a 2x-concentrated protease inhibitor cocktail (PIC). NC = negative control. **(E)** Luminescence in conditioned CCMM of inducible reporter strain cultures grown for 3 days on borosilicate glass beads, inoculated overnight with either *B. subtilis* or *E. coli,* and simultaneously supplemented with 20 mg/mL bovine serum albumin (BSA). Data from one experiment. Each data point represents one biological replicate as the average of two independent measurements. Horizontal bars represent means. **(F)** Data from panel E normalized by fungal colony dry weight (mg). Horizontal bars represent means. One-way ANOVA with Dunnett’s multiple comparisons test. ns = p > 0.05, *p< 0.05, **p < 0.01, ***p < 0.001, ****p < 0.0001. Horizontal bars indicate means.

*E. coli* has an inhibitory effect on *C. cinerea* growth, as opposed to *B. subtilis* [5]. Thus, we reasoned that smaller fungal colonies would secrete less cNluc and corrected for this effect by accounting for colony size. However, normalization by fungal colony dry weight did not affect data interpretation (Fig. 2B).

We then hypothesized that *E. coli* somehow negatively impacts cNluc stability or activity under our experimental conditions. To test this hypothesis, we grew the constitutive reporter strain axenically on liquid CCMM for 3 days and collected the conditioned medium, which by then should contain high amounts of secreted enzyme. Following filter-sterilization, the conditioned medium was aliquoted and either directly incubated at 28 °C (negative control) or first inoculated with *B. subtilis* or *E. coli*. After 12 hours of incubation, only medium inoculated with *B. subtilis* exhibited luminescence levels comparable to those registered at the start of the experiment. In the medium inoculated with *E. coli*, a 70% decrease in luminescence was observed, while luminescence was no longer detectable in the negative control (Fig. 2C). These results suggest that, contrary to our expectations, loss of cNluc activity occurs independently of *E. coli* in *C. cinerea* conditioned medium. Remarkably, *B. subtilis* prevents this phenomenon almost completely.

This unexpected finding prompted investigations of the potential mechanisms responsible for the loss of cNluc activity in *C. cinerea* conditioned medium and the protective effect of *B. subtilis*. Common causes of loss of enzymatic activity are inhibition, denaturation, and degradation of the enzyme. *C. cinerea* secretes proteases that may compromise cNluc integrity potentially leading to degradation [6]. On the other hand, *B. subtilis* is known to secrete high levels of extracellular proteins that could serve as alternative substrates for *C. cinerea* proteases [16, 17]. Therefore, we hypothesized that the *B. subtilis* secretome protects cNluc from degradation by *C. cinerea* proteases. If this were true, we could expect other protease substrates (e.g. exogenous proteins or peptides) to have a protective effect upon cNluc once added to the conditioned medium. Similarly, addition of protease inhibitors should also stabilize cNluc.

We repeated the experiment supplementing cNluc-containing medium with either bovine serum albumin (BSA), concentrated LB medium, or a protease inhibitor cocktail (PIC) (Fig. 2C-D). Consistent with our hypothesis, both BSA and LB showed remarkable protective effects upon cNluc. BSA in particular completely preserved luminescence levels over the 12-hour incubation period. On the other hand, the protective effect of PIC did not extend past 2 hours post inoculation (Fig. S4E), suggesting that PIC is unstable in our experimental conditions (e.g. duration, temperature, medium pH).

We next asked if we could harness the remarkable capacity of BSA to prevent cNluc degradation in our challenge assays. Specifically, we tested if the degradation of cNluc in the presence of *E. coli* can be prevented by BSA supplementation. However, *E. coli* still did not appear as the strongest inducer of *cclys1*p when BSA was added together with either *B. subtilis* or *E. coli* (Fig. 2E-F). One potential explanation for this is that BSA adds to the protective effect of the *B. subtilis* secretome, resulting in even higher levels of cNluc accumulation. Moreover, BSA is possibly consumed by both bacteria and *C. cinerea*, as its supplementation increased both fungal and bacterial biomass (Fig. S4C-D). Introducing an additional nutrient source like BSA into our system may alter fungal-bacterial interactions and, consequently, *cclys1* expression.

### Validation of a new cultivation set-up for the high-throughput screening of antibacterial defence elicitors in *C. cinerea* using the inducible cNluc reporter system

The main advantages of the cNluc reporter over previous approaches to monitoring *cclys1* expression (transcriptomics, qRT-PCR, *cclys1*p-*dTomato* reporter strain) are minimal hands-on time and quantifiability. We also found that the new reporter is more sensitive, as results can be obtained with lower amounts of fungal biomass compared to the other methods. For qRT-PCR or immunoblot analysis, for example, at least 3 fungal plates of 55 mm diameter need to be pulled into one biological replicate to extract sufficient amounts of total RNA or protein. On the other hand, a single 55 mm colony of cNluc reporter strain already produces measurable luminescence (Fig. 1C-D; Fig. 2A). These features make the system suitable for the simultaneous testing of different candidate inducers of antibacterial defence. Yet, our semi-liquid cultivation system is still relatively laborious in terms of setup time, space, and resources (e.g. medium volume), which ultimately limits the experiment size.

Thus, we tested the inducible cNluc reporter for antibacterial defence induction in a novel 96-well setup, which would be compatible with high-throughput screenings. For this purpose, oidia were inoculated into CCMM to a final OD_600_ of 1 and distributed in 100 µL aliquots across all wells. Following 7 days of incubation, each row was inoculated overnight with different bacteria (Fig. 3A), including *B. subtilis* and *E. coli* as positive controls. The other bacteria were selected from different ecological niches to assess whether *cclys1* induction is bacterium-or niche-specific. *B. megaterium* is a soil saprotroph like *B. subtilis*, while *Serratia marcescens* and *Pseudomonas syringae* are opportunistic human and plant pathogens, respectively [18–20]. *B. subtilis* 168, a lab-domesticated strain deficient in the production of specific secondary metabolites (e.g., antifungal lipopeptides), was included to test the role of these compounds in the induction of *cclys1* [21–23]. To ensure bacterial survival and growth in this set-up, axenic exponential cultures of the above bacteria were inoculated into a 96-well plate and their respective ODs_600_ measured following overnight incubation (Fig. S5A).

**Figure 3.**
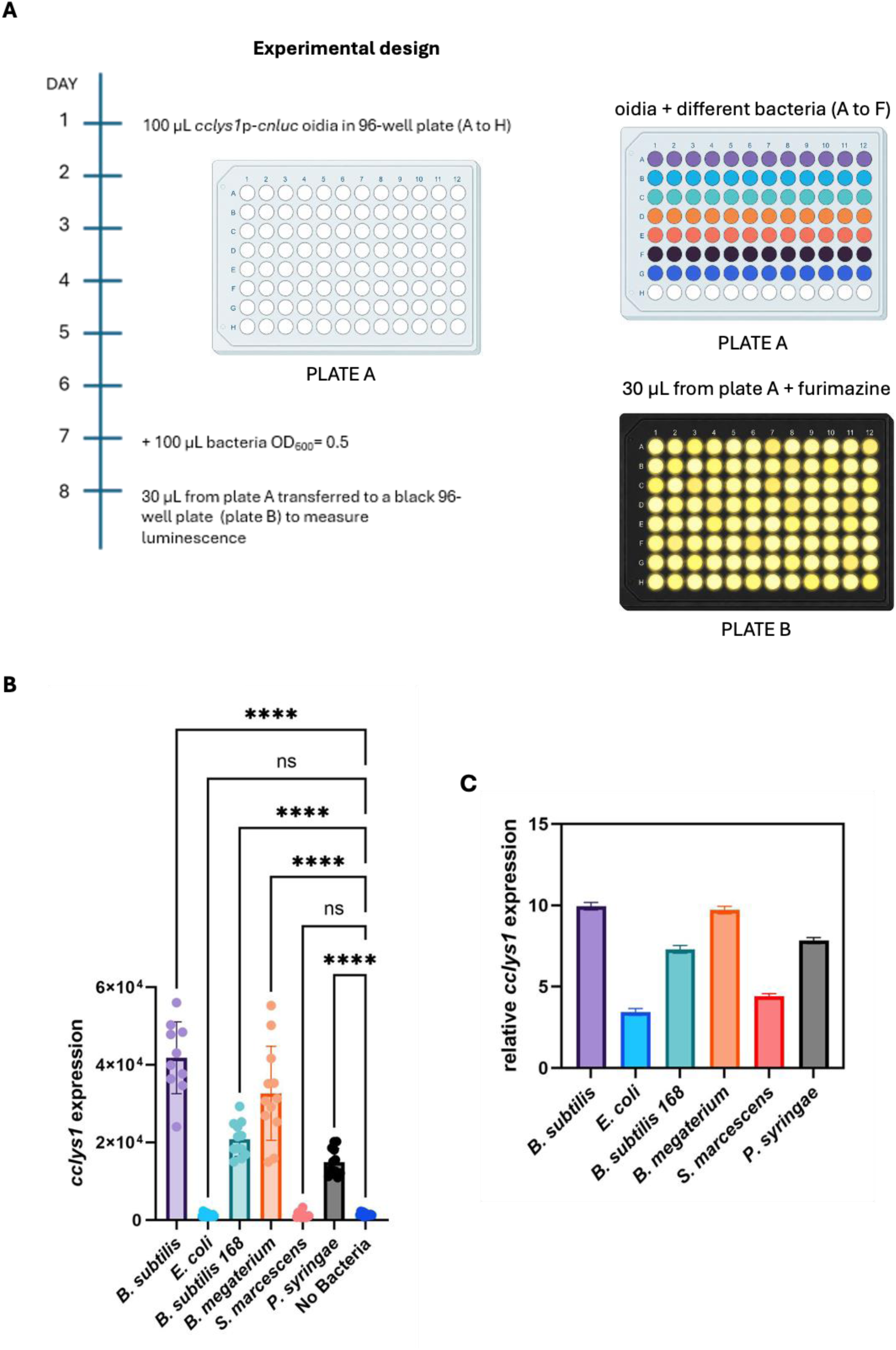
Validation of a new cultivation set-up for the high-throughput screening of antibacterial defence elicitors using the inducible reporter strain. **(A)** Experimental design for a high-throughput luminescence-based screening of *cclys1*p induction in a 96-well plate using germinated oidia from the inducible reporter strain. Each colour in plate A represents a different treatment. **(B)** Induction levels of *cclys1*p, grown as in panel A, in response to overnight inoculation with 6 different bacterial species, as determined by proxy of luminescence measured in the conditioned CCMM. Data from one experiment. Each data point corresponds to the luminescence measured in one well. The bar chart also shows means and standard deviations. One-way ANOVA with Dunnett’s multiple comparisons test. ns = p > 0.05, *p < 0.05, **p < 0.01, ***p < 0.001, ****p < 0.0001. **(C)** Treatment-specific expression of *cclys1* in the inducible reporter strain relative to the ‘No Bacteria’ control, grown as in panel A, as determined by qRT-PCR analysis with primers qPCR_*lys1*_FW1 and qPCR_*lys1*_RV1 (sequences provided in Table S2). Relative *cclys1* expression is reported as Log2 fold change. The bar chart shows the means and standard deviations (< 0.3) of 3 technical replicates.

Significant levels of luminescence, relative to the ‘No Bacteria’ negative control, were measured in all wells inoculated with *Bacillus* species and with *P. syringae (*Fig. 3B). qRT-PCR analysis confirmed comparable relative levels of expression (Fig. 3C and S5B). However, no luminescence was detected in response to either *E. coli* or *S. marcescens*. As qRT-PCR shows that *cclys1* is upregulated also upon these two treatments in this set-up, lack of detectable luminescence is probably again due to degradation of cNluc.

## Discussion

*Coprinopsis cinerea* responds to antagonistic bacteria with induction of genes mostly encoding secreted molecules [6]. Based on the experimentally proven antibacterial activity of many of these molecules, this response is interpreted as a specific antibacterial defence mechanism. Little is known about the elicitors of this response, which may range from physical contact to soluble molecules to abiotic stress factors like starvation caused by nutrient competition [6, 7]. To identify such elicitors, reporter systems, in which the coding region of a reporter protein is under the control of the promoter of an antibacterial defence gene, can be used [7].

In this work, we introduce a new reporter system, based on the secretion of the deep-sea shrimp luciferase Nluc, adapted for expression in *C. cinerea* by codon-optimization (cNluc). Compared to the intracellular accumulation of other reporter proteins (e.g., dTomato), cNluc ensures minimal hands-on time for sample processing, as enzymatic activity can be detected in small aliquots of fungal culture medium. The enzymatic activity of cNluc can also be quantified by measuring luminescence, which provides valuable information on the level of gene induction.

While luciferase-based reporter systems have been already established in filamentous fungi, the reporter protein accumulates intracellularly in all these instances [24, 25]. Expression of a firefly cytoplasmic luciferase (*LUC*) was achieved previously in *Aspergillus nidulans* and *Neurospora crassa*, in the latter for monitoring the activity of a circadian clock gene [24]. In contrast, cytoplasmic LUC is reportedly non-functional in *C. cinerea*, despite codon-optimization and 5’end intron addition [8, 26]. More recently, the original (non-secreted) Nluc variant was instead successfully employed in *C. cinerea* for the identification of novel endogenous promoters with strong expression [25]. Notably, cytoplasmic luciferase limits the experimental design to protoplasts or spore-originating fungal colonies grown in set-ups where luminescence can be measured directly [24, 25, 27]. In contrast, luciferase secretion ensures greater versatility across growing conditions.

To our knowledge, this is the first example of a non-fungal protein, without major modifications, successfully secreted by a filamentous fungus. Secretion of non-fungal proteins, including fluorescent reporters, is notoriously difficult in filamentous fungi [28, 29]. This limitation is often addressed by fusing the heterologous fluorescent protein to a secreted native protein [30, 31]. Here, we show that the signal peptide sequence of *cclys1* is sufficient to mediate secretion of functional cNluc in *C. cinerea*. At the same time, cNluc maintains all the advantages of the original Nluc luciferase over fluorescent proteins, which include smaller size, brighter signal, and lack of autofluorescence [32].

Another popular approach to building secretion-based reporters in filamentous fungi relies on the heterologous expression of fungal laccases [33]. Laccases are secreted, copper-containing enzymes that can oxidize the artificial substrate 2,2′-azino-di-(3-ethylbenzthiazolinsulfonate) (ABTS), forming a coloured (purple to green) product. Thus, laccase activity can also be measured in the fungal culture medium with colorimetric assays using ABTS. However, this approach is not suitable for gene expression analysis in *C. cinerea*, since the fungus already possesses 17 endogenous laccases, some of which are constitutively active [34, 35]. Expression of cNluc completely eliminates this issue.

Collectively, our data show that the inducible cNluc reporter produces readily quantifiable results. In addition, we found that the system is unexpectedly sensitive. Luminescence upon *cclys1* induction can still be detected when growing the reporter strain in a 96-well plate. The combination of these properties allows the high-throughput screening and comparison of candidate elicitors. However, we also observed that, under specific conditions, cNluc activity may be reduced or completely lost due to extracellular factors, such as *C. cinerea* secreted proteases, leading to false-negative results.

Degradation of cNluc by *C. cinerea* secreted proteases can be exacerbated by co-cultivation with certain bacteria. For example, *E. coli* causes the significant overexpression of at least one *C. cinerea* gene coding for a secreted cysteine-dependent endopeptidase (CC1G_05940T0) [6]. Bacteria like *S. marcescens*, which secretes high levels of proteases, contribute to the degradation of cNluc directly [36, 37]. We tried to address this issue by supplementing an excess of proteinaceous substrates. As expected, saturation of protease activity with BSA protects cNluc from digestion *in vitro*. Unfortunately, BSA is also a carbon and nitrogen source for both fungi and bacteria and cannot be supplemented to the challenge assays without significantly altering the physiology of the interacting organisms. For example, *E. coli* reached significantly higher ODs_600_ with BSA supplementation while resulting in comparable levels of luminescence. It is unclear whether BSA, by promoting *E. coli* growth, 1) relieves both nutrient and interference competition, resulting in lower levels of *cclys1* induction or 2) causes even higher levels of antibacterial defence induction, including overexpression of proteases, which may still lead to partial cNluc degradation. In the absence of an appropriate mitigation strategy, the system remains prone to false negatives.

Notably, the cNluc reporter faithfully recapitulates *cclys1* expression levels induced by *Bacillus* species. We believe that this is mostly due to the protective effect of *Bacillus* abundant secretome against cNluc digestion by *C. cinerea* secreted proteases. Moreover, *Bacillus subtilis* does not strongly boost protease expression in *C. cinerea* [6]. This makes the cNluc reporter particularly suitable for the screening of *Bacillus* knock-out strain collections, such as the one available in the *B. subtilis 168* background (https://www.addgene.org/kits/grosslab-bsubtilis-collections/)).

Based on these data, we conclude that the cNluc reporter system is suitable for monitoring the induction of a transcriptional response in *C. cinerea* and most likely other filamentous fungi across a wide range of experimental set-ups, including one compatible with high-throughput screenings. However, negative results should always be validated by qRT-PCR analysis. To make this reporter more robust, future work could focus on engineering a protease-stable version of cNluc.

## Supplementary Figures

**Figure S1.**
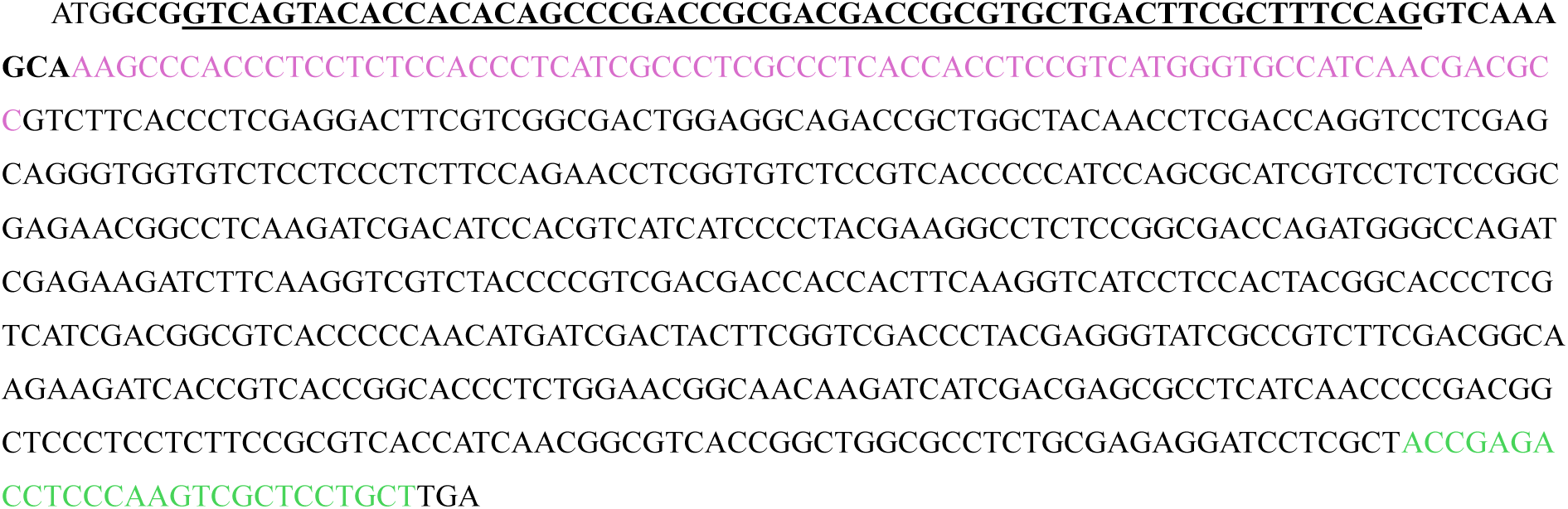
The *cnluc* coding sequence. The original Nanoluc (Nluc) coding sequence was codon-optimized for expression in *C. cinerea* and modified to C-terminally tag the protein with the 1D4 epitope of bovine rhodopsin (Rho1D4, highlighted in green). A sequence coding for the secretion signal of *C. cinerea lys1* (*cclys1*, CC1G_03076T0) was added to the 5’end of the Nluc-Rho1D4 coding sequence (highlighted in purple). To further increase expression efficiency, an intron-containing region from the *P. chrysosporium mnp* gene (in bold, with intron underlined) was introduced between the start codon and the signal peptide sequence. Intron splicing leaves behind 4 amino acid-coding triplets at the 3’end of the sequence. This modified *Nluc* sequence encodes the novel reporter protein cNluc.

**Figure S2.**
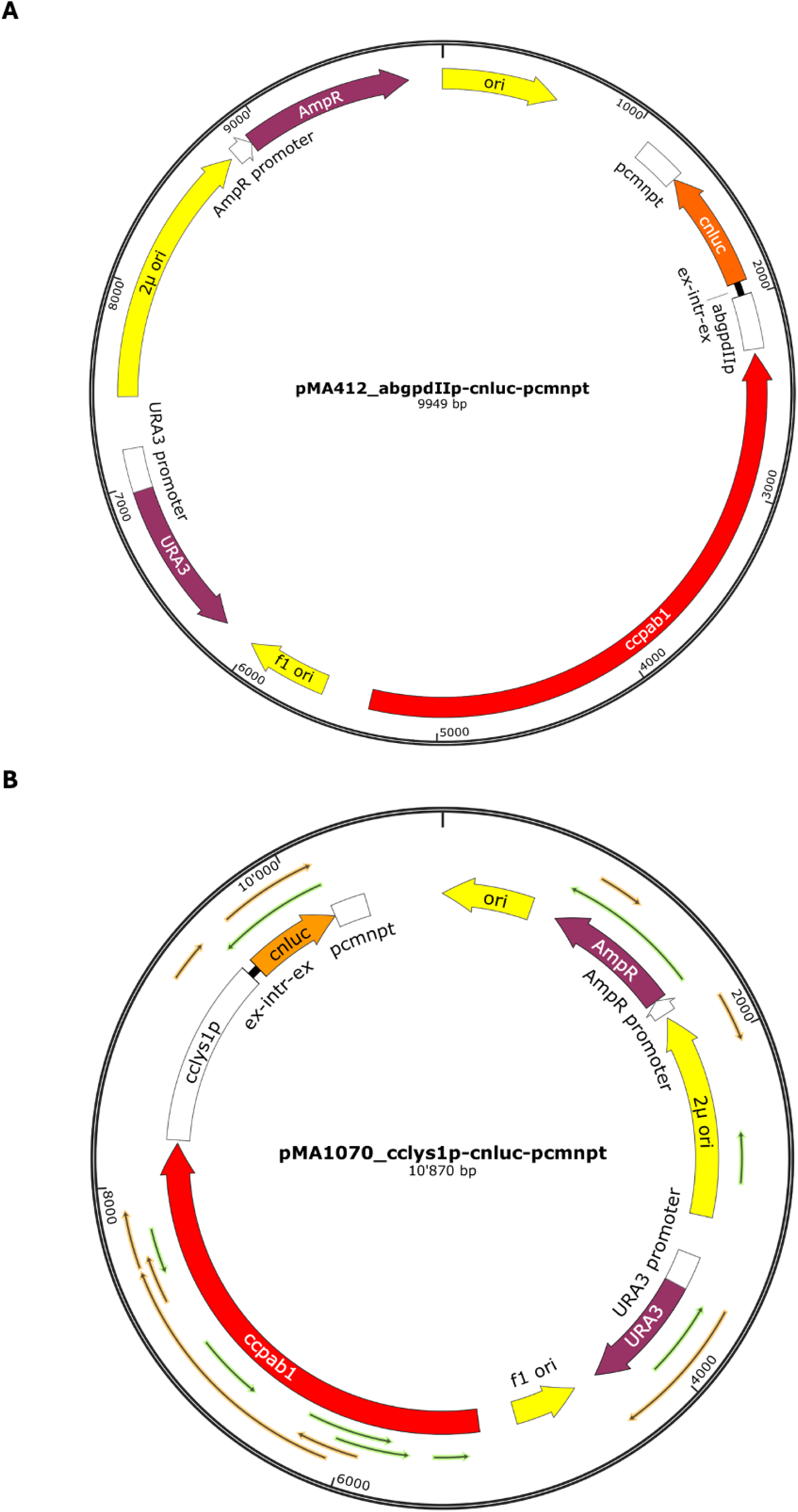
Plasmids used to generate *C. cinerea* constitutive and inducible cNluc reporter strains. **(A)** Plasmid used to generate the constitutive cNluc reporter strain. Here, the *cnluc* gene (orange) is under the control of the gpdII promoter from *A. bisporus* (*AbgpdII*p). **(B)** Plasmid used to generate the inducible cNluc reporter strain. The cNluc gene is instead under the control of the *lys1* promoter from *C. cinerea* (*cclys1*p). In both plasmids, transcription of *cnluc* is under the control of the *mnp* terminator region of *P. chrysosporium* (*pcmnp*t). Moreover, an intron-containing region from *P. chrysosporium gpd* (ex-intr-ex) was introduced between the promoter sequence and the *cnluc*. Both plasmids also feature an Ampicillin resistance cassette (*AmpR*) for selection in *E. coli* and a functional *pab1* gene from *C. cinerea* strain Okayama 7 (*ccpab1*) for selection in *C. cinerea*.

**Figure S3.**
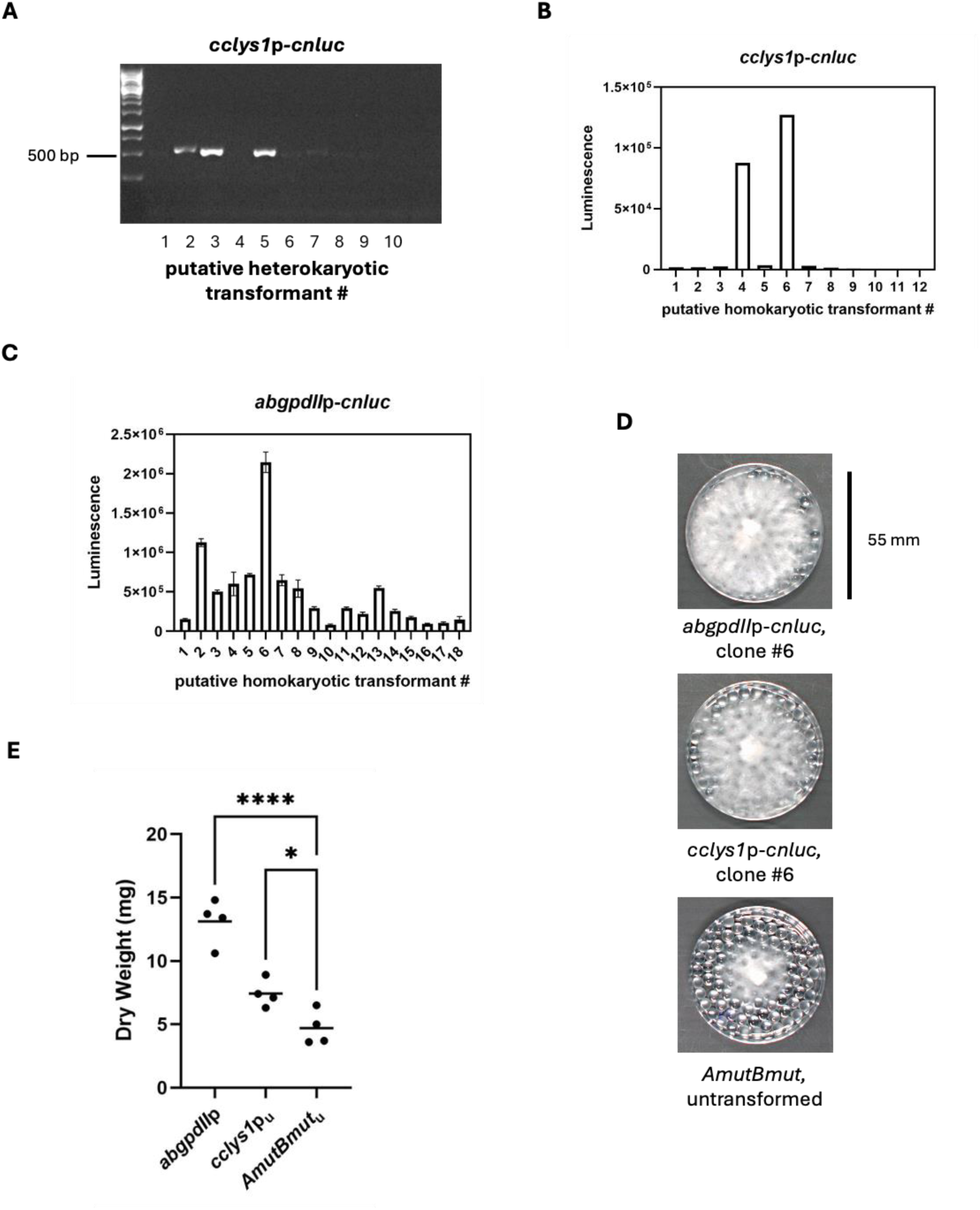
Selection of stable homokaryotic *C. cinerea* transformant strains. **(A)** Colony PCR screening of putative inducible cNluc transformants (*cclys1*p-*cnluc*) with primers targeting the *cnluc* gene (primer sequences in Table S2). Expected amplicon size = 556 bp. **(B)** Luminescence of conditioned CCMM from putative inducible cNluc homokaryotic transformants (*cclys1*p-*cnluc*) after 3-day cultivation on borosilicate glass beads and overnight inoculation with *E. coli* CFS. Putative homokaryons were germinated from single oidia of PCR-confirmed heterokaryotic transformant #5 (in panel A). Bars represent the means of two independent measurements. **(C)** Luminescence of conditioned CCMM from cultures of putative constitutive cNluc homokaryotic transformants (*abgpdII*p-*cnluc*). Transformants were grown for 3 days on borosilicate glass beads. Putative homokaryons were germinated from single oidia of a putative heterokayotic transformant identified on selective medium (data not shown). The bar chart shows the means of two independent measurements with standard deviations. **(D)** Representative images of confirmed homokaryotic cNluc reporter strains grown axenically for 3 days in CCMM on borosilicate glass beads: constitutive, clone #6 (*abgpdII*p-*cnluc*), inducible, clone #6 (*cclys1*p-*cnluc*), and untransformed *C. cinerea AmutBmut*. **(E)** Individual colony dry weight (mg) of cNluc reporter strains grown axenically in CCMM on glass beads for 3 days: constitutive (*abgpdII*p), inducible unchallenged (*cclys1*p_u_), and untransformed *AmutBmut* (*AmutBmut*_u_). Each data point represents a biological replicate. Horizontal bars indicate means. One-way ANOVA with Dunnett’s multiple comparisons test. ns = p > 0.05, *p< 0.05, **p < 0.01, ***p < 0.001, ****p < 0.0001.

**Figure S4.**
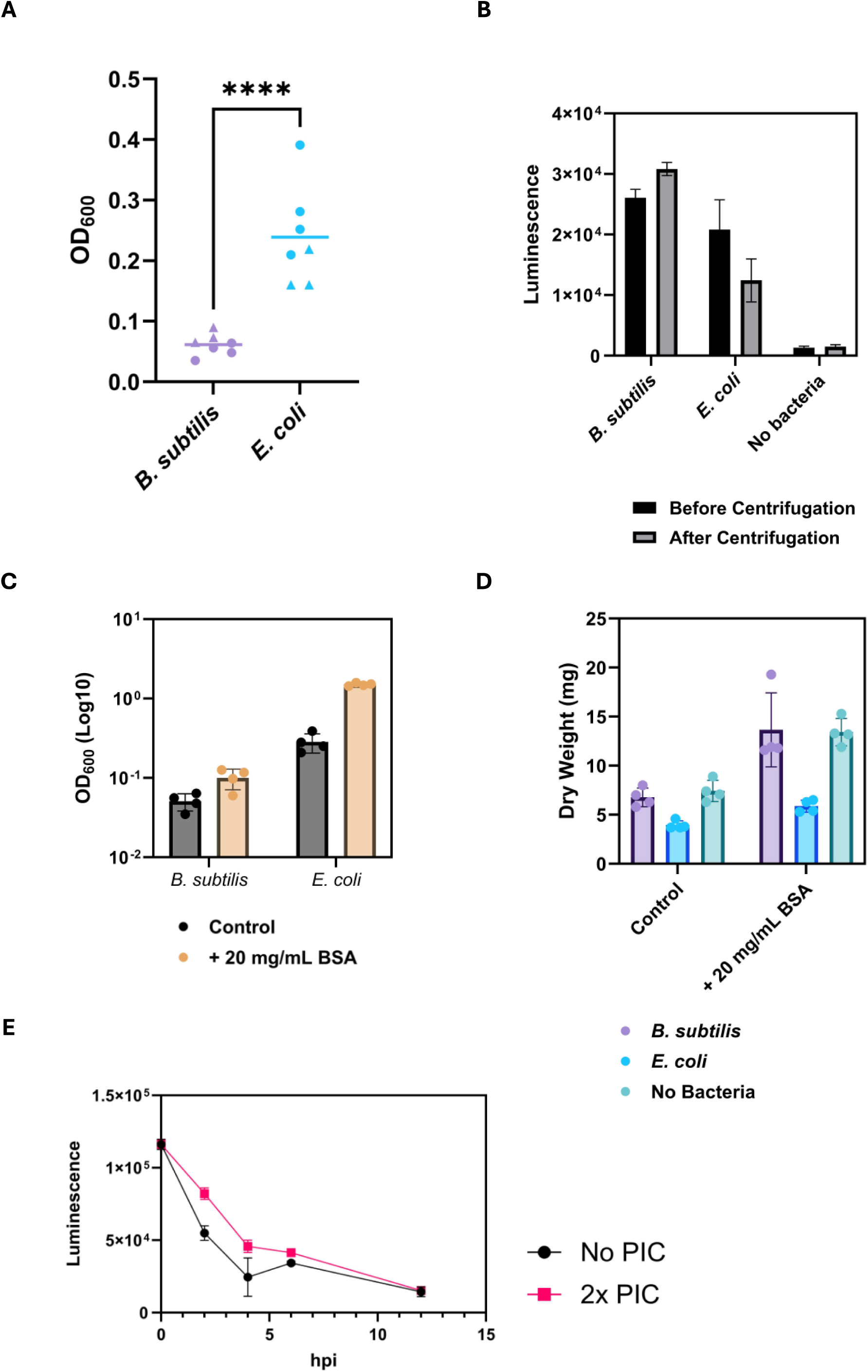
Validation of the inducible reporter strain for quantification of antibacterial defence induction. **(A)** Optical density at 600 nm (OD_600_) of *B. subtilis* and *E. coli* cultivated overnight with the inducible cNluc reporter strain in 5 mL CCMM. Data from two experiments, each represented by a different symbol. Horizontal bars indicate means. Unpaired T-test. *p < 0.05, **p < 0.01, ***p < 0.001, ****p < 0.0001. **(B)** Luminescence in conditioned CCMM of inducible cNluc reporter colonies grown for 3 day on glass beads, either axenically (no bacteria) or inoculated overnight with *B. subtilis* or *E. coli*. Measurements were taken before and after centrifugation. Data from one experiment. Bars represent mean measurements with respective standard deviations. **(C)** OD_600_ of *B. subtilis* and *E. coli* cultivated overnight with the inducible cNluc reporter strain in 5 mL CCMM, with or without BSA. Data from one experiment. Bars represent means with respective standard deviations. **(D)** Individual colony dry weight (mg) of inducible cNluc reporter colonies grown for 3 days in CCMM on glass beads, either axenically or overnight with *B. subtilis* or *E. coli*, with or without BSA. Bars indicate means with respective standard deviations. **(E)** Luminescence of sterile cNluc-containing CCMM over 12 hours (hpi = hours post inoculation), with or without supplementation with a 2x-concentrated protease inhibitor cocktail (PIC). Each data point represents the average of three technical replicates. Error bars show standard deviations between technical replicates.

**Figure S5.**
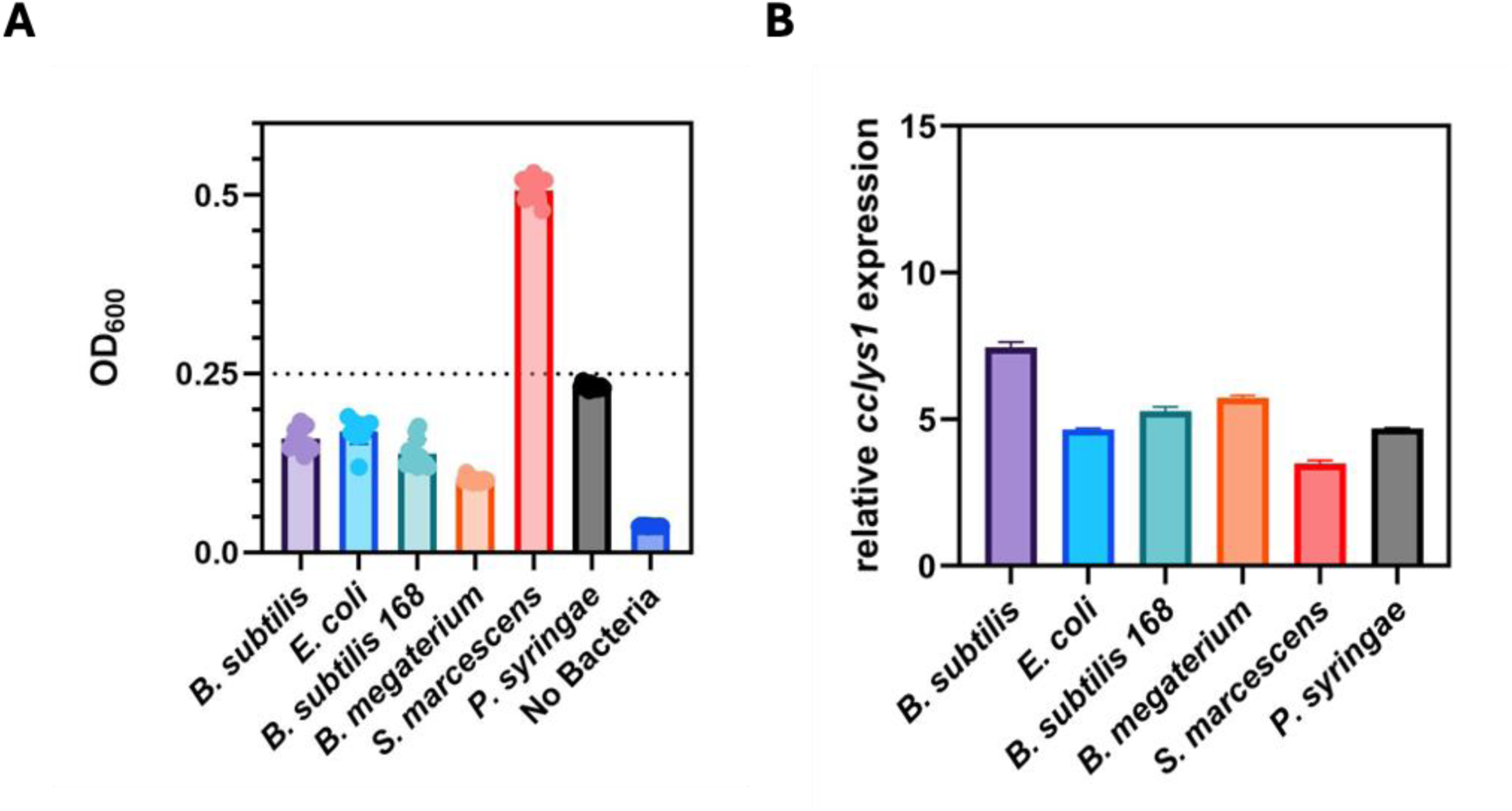
Validation of a new cultivation set-up for the high-throughput screening of antibacterial defence elicitors using the inducible reporter strain. **(A)** Optical density at 600 nm (OD_600_) of different bacteria grown axenically overnight from starting OD_600_ = 0.25 in CCMM in a 96-well plate. Data from one experiment. Each data point corresponds to the OD_600_ in 1 well. Bars show means. **(B)** Treatment-specific expression of *cclys1* in the inducible cNluc reporter strain relative to the ‘no bacteria’ control, as determined by qRT-PCR analysis using primers qPCR_*lys1*_FW2 and qPCR_*lys1*_RV2 (sequences in Table S2). Relative *cclys1* expression is reported as Log2 fold change. The bar chart shows the means and standard deviations (< 0.3) of 3 technical replicates. The inducible cNluc reporter strains and the bacteria were grown as described in Fig. 3 and in ‘Materials and Methods’.

## Supplementary Tables

**Table S1.** List of fungal (orange) and bacterial (light blue) strains used in this study.

| Species | Strain | Source |
| --- | --- | --- |
| *Coprinopsis cinerea* | *AmutBmut* | Own Lab |
| *Saccharomyces cerevisiae* | *W303 MATa* | Own Lab |
| *Escherichia coli* | *DH5α* | Own Lab |
| *Escherichia coli* | *Nissle 1917* | W. D. Hardt |
| *Bacillus subtilis* | *NCIB 3610* | R. Losick |
| *Bacillus subtilis* | *168 Trp<sup>+</sup>* | U. Sauer |
| *Bacillus megaterium* | *DSM90* | DSMZ |
| *Serratia marcescens* | unknown | P. Lüthy |
| *Pseudomonas syringae* (pv. tomato) | *DC3000* | J. Vorholt |

**Table S2.**
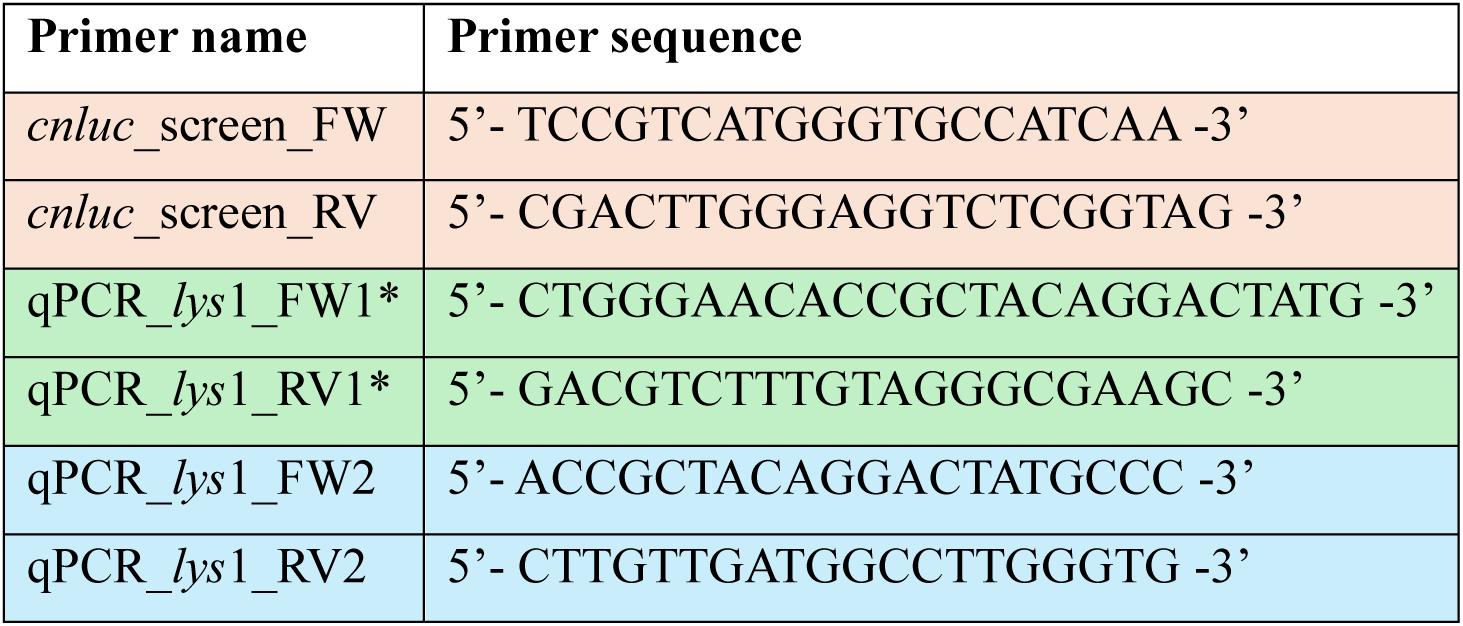
List of primers used in this study. Paired primers share the same colour. The primer pair designated with an asterisk (*) has been published previously [15].

## Notes

### Competing Interest Statement

The authors have declared no competing interest.

### Funding

- Swiss National Science Foundation: 310030-212435
- ETH Zurich

